# Visual and auditory information shed light on savings mechanisms

**DOI:** 10.1101/569186

**Authors:** Olivier White, Marie Barbiero, Quentin Maréchal, Jean-Jacques Orban de Xivry

## Abstract

Successful completion of natural motor actions relies on feedback information delivered through different modalities, including vision and audition. The nervous system weights these sensory inflows according to the context and they contribute to the calibration and maintenance of internal models. Surprisingly, the influence of auditory feedback on the control of fine motor actions has only been scarcely investigated alone or together with visual feedback. Here, we tested how 46 participants learned a reaching task when they were provided with either visual, auditory or both feedback about terminal error. In the VA condition, participant received visual (V) feedback during learning and auditory (A) feedback during relearning. The AV group received the opposite treatment. A third group received visual and auditory feedback in both learning periods. Our experimental design allowed us to assess how learning with one modality transferred to relearning in another modality. We found that adaptation was high in the visual modality both during learning and relearning. It was absent in the learning period under the auditory modality but present in the relearning period (learning period was with visual feedback). An additional experiment suggests that transfer of adaptation between visual and auditory modalities occurs through a memory of the learned reaching direction that acts as an attractor for the reaching direction, and not via error-based mechanisms or an explicit strategy. This memory of the learned reaching direction allowed the participants to learn a task that they could not learn otherwise independently of any memory of errors or explicit strategy.

## Introduction

The brain relies on sensory feedback to correct inaccurate movements and improve the outcome of subsequent actions. The body is equipped with different biological sensors that serve this purpose. For instance, vision (Held and Freedman, 1963) and proprioception (Wolpert et al., 1995; Sarlegna and Sainburg, 2009) are known to yield critical and complementary information (van Beers et al., 2002). Surprisingly, very little attention has been paid to audition in this respect despite the fact this inflow is the main sensory feedback modality for speech movements. Humans modulate the loudness of their speech in function of the loudness of the environment and in function of how they perceive their own voice (Lane and Tranel, 1971). Similarly, artificially altering the auditory feedback results in a modification of the vocal output (Chen et al., 2007). If such modifications are maintained consistently for several exposures, then, it leads to a long-term change in speech production (Bomze et al., 1998).

Both visual and auditory feedback modalities can be used to guide upper limb and speech movements. For instance, visual information about one’s own speech movements can improve language training (Katz and Mehta, 2015). Similarly, the ability to use auditory feedback to maintain optimal motor output extends beyond speech production. When sound is not consistent with an action (e.g. delayed auditory feedback), it may interfere the smooth course of an action and even result in destructive interactions. Altering auditory feedback during the performance of sequential motor actions such as playing the piano interferes with the production of the melody (Pfordresher, 2005, 2012; Furuya and Soechting, 2010). Audio-motor interactions are numerous while playing musical instruments (Zatorre et al., 2007; Cao et al., 2016) or, more generally, in other types of tasks (Sigrist et al., 2013). Proper use of auditory information can also result in constructive interaction. For instance, providing auditory feedback about motor performance is sufficient to induce reliable learning in a natural gait learning task (Zanotto et al., 2013) or in a trunk-arm rowing task (Sigrist et al., 2014) and also in a classical experimental cursor motion tracking paradigm (Rosati et al., 2012). Auditory feedback can be used for sensorimotor adaptation of reaching movements even though such adaptation mainly relies on visual and proprioceptive information. For instance, changing the location and/or loudness of the sound allows humans to derive the motor error and to adapt to a perturbation (Oscari et al., 2012; Schmitz and Bock, 2014). This suggests that sensory substitution of visual information by auditory information is sufficient to induce motor adaptation. But, are auditory and visual information treated equally by the brain or do they provide complementary information?

Our main interest was to study how learning with one modality (e.g. visual) could be transferred to relearning in another modality. Savings correspond to the ability to learn faster on the second exposure during a learning task as compared to the first exposure. This well-known phenomenon has been observed in a wide range of declarative (e.g. Ebb¡nghaus, 2014) and motor learning tasks (Brashers-krug et al., 1996; Krakauer et al., 2005). In motor adaptation tasks, several mechanisms have been proposed to account for savings. Some authors suggested that savings could arise from a memory of errors (Herzfeld et al., 2014), from the reliance on previous motor memories (Smith et al., 2006), from the use of an explicit strategy (Haith et al., 2015; Morehead et al., 2015) and/or from a reward-based memory of successful motor actions (Huang et al., 2011; Shmuelof et al., 2012; Orban de Xivry and Lefèvre, 2015) as well. Therefore, the controversy around these mechanisms remains open.

While experimental manipulations exist to minimize the contribution of the explicit strategy (Taylor et al., 2014; Morehead et al., 2015) or the reliance on previous motor memories (Huang et al., 2011; Orban de Xivry and Lefèvre, 2015), reducing the contribution of memory of errors mechanisms has proven more difficult. Here, we reasoned that if savings was observed across modalities, it could not be due to a memory of errors given their different modalities. In a first experiment, we tested the transfer of motor adaptation between visual and auditory sensory modalities. In a second experiment, we tested whether the observed savings is due to an explicit strategy or to a memory of the successful motor actions learned during the first exposure (Huang et al., 2011; Orban de Xivry and Lefèvre, 2015).

## Materials and Methods

### Participants

Forty-six right-handed adults were included in this study. Experiment 1 comprised 34 participants (17 females, 24.3 ± 3.9 years old) and Experiment 2 involved 12 different participants (8 females, 2 ± 2.2 years old). All participants were volunteers, healthy, without neuromuscular disease and with normal or corrected to normal vision. Each participant had no auditive impairment and was musically trained. The experimental protocol was carried out in accordance with the Declaration of Helsinki (1964) and the procedures were approved by the local ethics committee of Université de Bourgogne-Franche Comté. All participants were naïve as to the purpose of the experiments and were debriefed after the experimental session.

### Apparatus and stimuli

Participants were comfortably seated in front of a virtual environment equipment with the head on a chin rest in a dimly illuminated room. They looked into two mirrors that were mounted at 90 degrees to each other, such that they viewed one LCD screen with the right eye and one LCD screen with the left eye. This stereo display was calibrated such that the physical locations of the robotic arms were consistent with the visual disparity information. Participants made 12-cm movements while holding on to a robotic device with the right hand (Phantom 3.0, SensAble Technologies, USA). Movements were performed in the natural reaching space in an upward-forward direction, involving shoulder and elbow movements, with the elbow pointing downwards.

### Experimental procedure

Participants performed shooting movement toward one of 5 green targets positioned on a circle (radius=12cm) at five different eccentricities (−40deg, −20deg, 0deg, 20deg, 40deg). The 0-deg direction corresponded to the vertical upward direction (Figure 1A). The 3d positions of the robot handle were mapped in real time to a grey cursor (diameter=3mm). A trial started when the cursor was positioned inside a starting green sphere (diameter=6mm). Then, as soon as one of the five targets appeared (diameter=6mm), the participant performed a rapid shooting movement through the target. Continuous visual feedback of the cursor trajectory was not provided. The trial ended when the radial distance from the starting position and the cursor was larger than 12cm. We instructed and trained participants to reach the target within around 250ms. After each trial, the target turned red or blue if movement times were longer than 300ms or shorter than 200ms, respectively.

**Figure 1:**
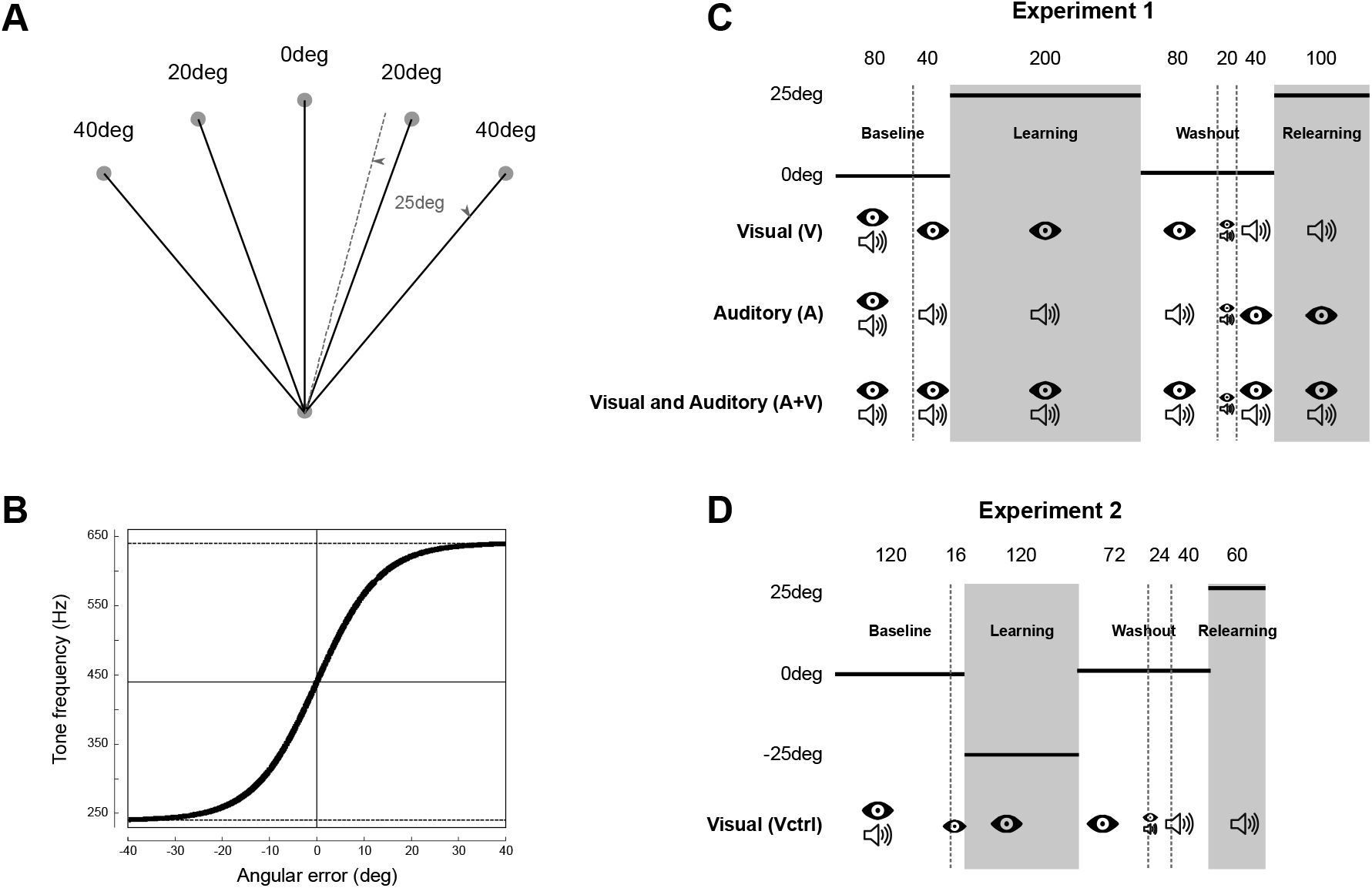
Illustration of the experimental procedures. **A)** Scaled trajectories of reaching movements toward the 5 targets in Experiment 1 (top circles). The dotted line represents a successful shooting movement toward the 40-deg target after full adaptation to a +25deg rotation in hand space. **B)** Pitch of the feedback sound (y-axis) in function of the angular error (x-axis). Resolution is optimized around 0-error (largest slope). The two horizontal dashed lines represent the saturation pitches (240Hz and 640Hz) and the horizontal solid line is positioned at the reference frequency of 440Hz. The vertical cursor is positioned at the angular error of 0deg. **C)** In Experiment 1, each group (rows) received different feedback modalities but were exposed to the same perturbation schedule. An “eye” icon and a “speaker” icon correspond to the presence of visual (V), auditory (A) or both (V+A) feedback information. The numbers on top represent the number of trials for each period, separated by dashed vertical and by the grey rectangles. **D)** Experimental schedule for Experiment 2.

After 500ms, a feedback about shooting error was provided through two different modalities. In the visual modality, a yellow cursor (diameter=6mm) was flashed during one second at the position the closest to the target. The error was therefore signed; the yellow cursor could appear on the left or on the right of the target. In the auditory modality, error information was provided through a sound defined the following way. A continuous sound was emitted for one second but with specific frequencies. During the first 500ms, its pitch followed a sigmoid function defined by:

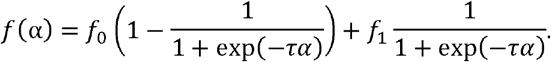

Pitch saturated at frequency f_0_ or f_1_ for angular errors α larger than −30deg or +30deg, respectively (Fig. 1B). The inflection point and largest slope of the sigmoid function was set at 0deg error in order to maximize the resolution of the auditory feedback around target position. The parameters were set as follows: f_0_=240Hz, f_1_=640Hz and τ=0.15deg^−1^. During the second 500ms, a single frequency tone of 440Hz that corresponded to a “*F*” in the main octave was played back. This tone provided a reference sound for a perfect shooting movement (error α=0deg). Participants interpreted the sign of the error by comparing the trial and reference pitches, these aptitudes being well developed in humans (Plack and Oxenham, 2005). Trial frequencies lower (resp. upper) than 440Hz were interpreted as angular errors to the left (resp. right) of the target and no error resulted in no identified differences between pitches. The 500-ms modulated by the angular error was emitted immediately after the trial in order to reinforce the link between the movement just performed and its auditory consequences. After the trial ended, the robot moved the hand back onto the starting circle. This sped up trials and prevented participants to perform active return movements.

In Experiment 1, participants were randomly dispatched in one of three groups, depending on feedback modality. In the “VISUAL” group (V→A, n=12), endpoint error was provided through visual modality and then through auditory modality. It was the opposite in the “AUDITORY” group (A→V, n=11). A third group was provided with both modalities for all trials (“AUDITORY *and* VISUAL”, V+A, n=11).

After briefing, participants were familiarized with a practice block of 20 trials to a single target at −15deg that were not considered for analysis. Cursor trajectories were not displayed online and feedback was provided both in the visual and auditory modalities. During the experimental session, participants performed 560 trials in the following order (Fig. 1C). The baseline block consisted of 120 reaching movements (24 to each target) to randomly selected targets (Fig. 1C, “baseline” sequence). During the first 80 trials, feedback was visual and auditory whereas it was either visual, auditory or both, depending on the group, for the last 40 trials. Then a visuomotor perturbation was introduced for the next 200 shooting movements (40 trials x 5 targets). The perturbation consisted of a +25deg clockwise rotation of the cursor for all groups. However, groups relied on different feedback to adapt their shooting movements (Fig. 1C, “learning” sequence). During the perturbed trials, participants had to move the hand 25deg to the left of the target in order to perform a successful trial (Fig. 1A, dotted lines). After this perturbation block, 140 trials were again performed in the same condition as baseline (Fig. 1C, “washout” sequence), without visuomotor rotation. During the first 80 trials, only feedback modality of that group was provided, like in the previous sequence. Next, participants could use both feedback modalities to adjust their movement in a short block of 20 trials. Finally, during the last 40 trials of this “washout” sequence, participants switched to the other feedback modality (i.e. “auditory” for the “VISUAL” group and vice-versa). The “*VISUAL and AUDITORY*” group had always both feedback modalities. In the last sequence (Fig. 1C, “re-learning”), the same perturbation was reintroduced in the other modality for 100 trials.

In Experiment 2 (n=12), we specifically tested for the contribution of an explicit strategy or of a reward-based memory of the learned movement direction. The procedure was similar to the “*VISUAL*” group in Experiment 1 and differed only for target positions and number of presentations (Fig. 1D). The baseline block consisted of 120 reaching movements to one of four targets (−90deg, −40deg, 10deg or 60deg)weon the basis of (Fig. 1C, “baseline” sequence). In the “learning” sequence, only three targets were presented (−90deg, −40deg and 10deg) and hand cursors were rotated 25deg counterclockwise (−25deg). In order to avoid overlearning, block design ensured that the same number of trials per target were presented in this condition (40 trials per target x 3 targets = 120 trials). The “washout” sequence consisted in 136 shooting movements to the same four targets as in the baseline sequence without visuomotor rotation. Finally, re-learning was tested in the “auditory modality but with different targets (−40deg, 10deg and 60deg) and with the opposite perturbation (clockwise, 25deg). The new set of targets was arranged so that the same hand trajectories (−65deg, −25deg and 15deg) as during the learning led to success. In other words, this experimental design allowed us to test savings. In hand space, the solution to the second perturbation of three targets ([−40deg, 10deg, 60deg]−25deg = [−65deg, −15deg, 35deg]; “re-learning”) was identical to the solution of the first perturbation ([−90deg, −40deg, 10deg]+25deg = [−65deg, −15deg, 35deg], “learning”).

### Data processing

Positions were recorded with a sampling rate of 500Hz. Movement start was detected when movement velocity exceeded 3cm/s for at least 100ms. Angular error of each movement was defined as the angular deviation at the end of the movement from the straight direction towards the target. In Experiment 1, we quantified, for each participant, the amount of learning, washout and relearning by looking at the difference between the mean of the angular errors during the first 5 trials and the mean of the angular errors during the last 15 trials in each sequence. We adopted the same procedure for Experiment 2 to account for the different number of targets among the three sequences. Namely, learning and relearning (using three targets) were assessed using the first 3 and last 9 trials while washout (using four targets) was quantified by comparing the difference in angular errors between the first 4 trials and the last 12 trials in that sequence.

The existence of savings was assessed by quantifying the speed of learning. To do so, we fitted an exponential function of the form *a*_1_*exp*(−*a*_2_*t*) + *a*_3_ to the angular error, where a_1_, a_2_ and a_3_ are constants that are fitted to the data. These three gains quantify the amplitude of adaptation, the adaptation rate and the value toward which the error converge, respectively. Sample size was fixed at 10+ subjects per group before the start of the experiments. This number was deliberately chosen higher than in previous experiments on savings, as we looked at a between-subject effects. We also report partial eta-squared for significant results to account for effect size. Quantile-quantile plots were used to assess normality of the data. Independent t-tests were conducted to compare data between groups (of unequal size) and paired t-tests were adopted to compare different conditions within groups. Data processing and statistical analyses were done using Matlab (The Mathworks, Chicago, IL).

## Results

In this experiment, we test the effect of a visual vs. auditory feedback on the adaptation to a visuomotor rotation and on the transfer of learning between these two modalities. We reasoned that such transfer would highlight the role of non-error-based mechanisms in savings because error-based saving mechanisms should depend on the modality of the error feedback. Participants from the V→A group received visual feedback during learning and auditory feedback during relearning. The second group (A→V) received auditory feedback during learning and visual feedback during relearning. Finally, we expected maximal adaptation for the participants who received both visual and auditory feedback during the entire experiment (V+A group). Except for the modality of the error feedback, the participants from these three groups were exposed to an identical schedule of visuomotor perturbations (Fig. 1C).

### Inability to adapt under auditory error feedback

After baseline trials, each group was confronted to a 25-deg visuomotor rotation (Fig. 2). Initially, the perturbation caused identical initial angular errors in the three groups (main effect of group on initial angular error: F_2,31_=1.74, p=0.19, 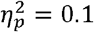). However, adaptation to the perturbation was absent in the group learning under auditory feedback (A→V) while the two other groups (V→A and V + A) could adapt normally to the perturbation (Learning period in Fig. 2A). The absence of learning under auditory feedback about terminal errors resulted in a larger final error for the A→V group compared to the two other groups (Fig. 2, second column), yielding a significant effect of group on the final error measure F_2,31_=56.25, p<10^−10^, 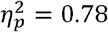). A Tukey post-hoc test to compare the final error across groups revealed that the group that learned in the AUDITORY modality (A→V) was different from the two other groups (A→V vs. V→A: p<10^−8^ and A→V vs. V+A: p<10^−8^). In other words, participants from the A→V group did not decrease their angular error as much as the two other groups (interaction between group and time, initial vs. late error: F_1,31_=58.17, p<10^−7^, 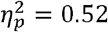). Indeed, late angular errors were significantly smaller than initial errors in the V→A group (t-test: t_11_=11.9, p<10^−6^) and V+A group (t_10_=9.31, p<10^−5^) but not in the A→V group (t_10_=1.05, p=0.318) that received auditory feedback only during the learning period.

**Figure 2.**
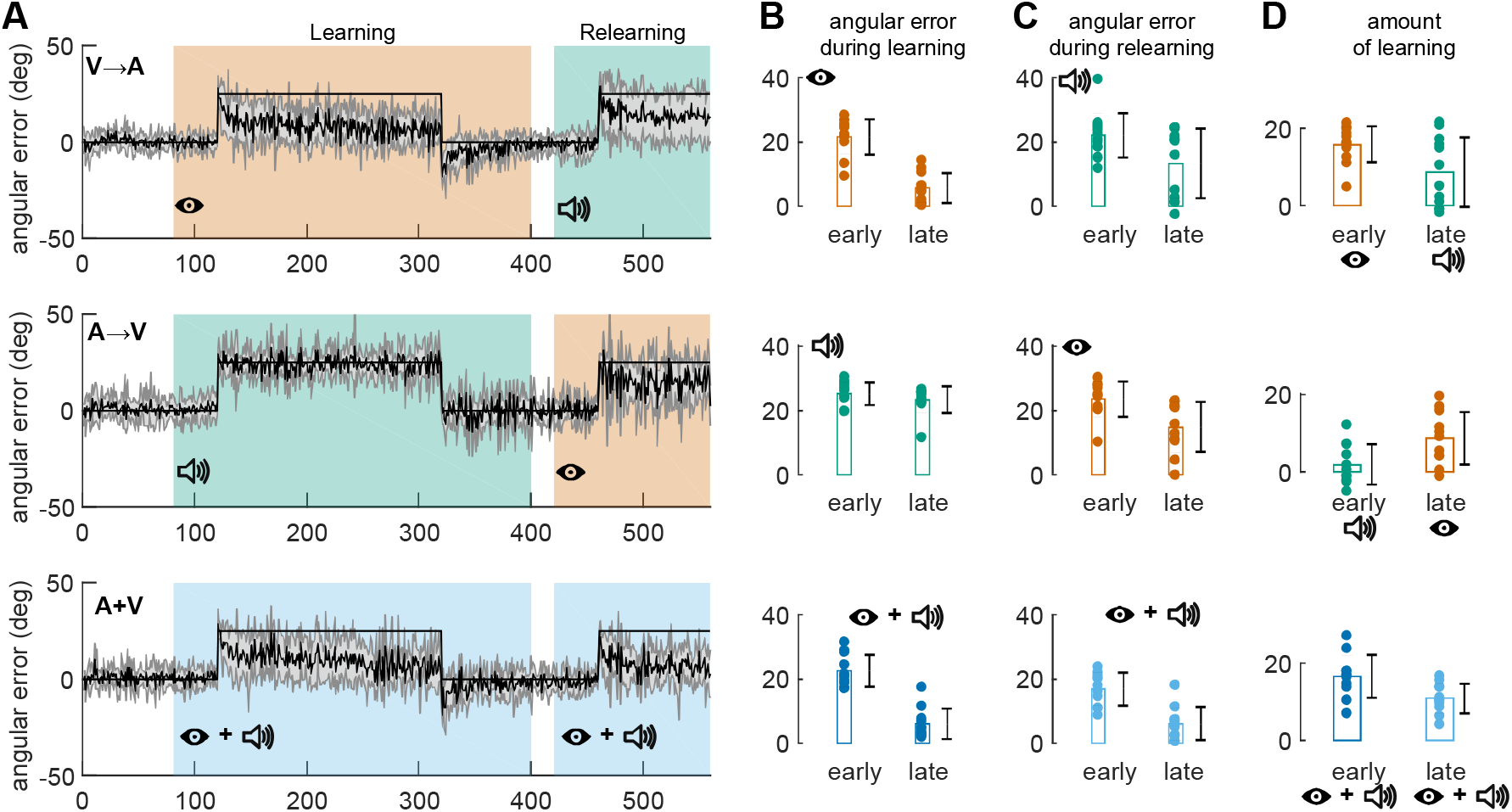
Evolution of the angular error over trials for the three groups of Exp. 1 (First line: V→A, second line: V→A, third line: V+A). **A)** Evolution of the angular error over the course of the experiment. Data are averaged across participants (solid line) and grey shaded area correspond to the standard error of the mean. Thin black line corresponds to cursor rotation. Green background: auditory feedback. Orange background: Visual feedback. Blue background: auditory and visual feedback. **B)** Average angular error early and late during the learning period. Early=first cycle (five trials); Late=last six cycles (30 trials). **C)** Average angular error early and late during relearning. **D)** Average amount of learning during the learning (early) and relearning (late) periods. For **B, C and D**, colour code is similar to A. On each plot, individual values are represented by dots for each measure. Rectangles represent the mean. Black error bars represent mean ± SE.

The lack of adaptation of the A→V group was confirmed by the limited after-effect for this group in the ensuing washout period under the same feedback modality (mean ± SD: −2.5±3.8deg) while the two other groups demonstrated a much larger after-effect (V→A: −13.57±3.6deg and V+A: −11.4±5.9deg). The after-effect of the A→V group was significantly smaller than in the other two groups (main effect of group: F_2,31_=19.03, p<10^−5^, 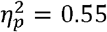; post-hoc Tukey test: A-V vs V-A: p<10^−5^; A-V vs A+V: p=0.0002 and V-A vs V+A: p=0.49).

Participants from all groups then performed 20 trials with concurrent visual and auditory feedback without perturbation (Fig. 1C), which were followed by 40 unperturbed shooting movements but with only the other feedback modality. In other words, the V→A and A→V groups switched to the other feedback modality (the V+A group still received V+A feedback). By the end of this washout phase, all groups performed reaching movements without significant errors (V→A: t_11_=1.13, p=0.28; A→V: t_10_=0.76, p=0.463; V+A: t_10_=0.57, p=0.583).

### Relearning under auditory feedback is possible after learning under visual feedback

In the relearning period, participants from the V→A and A→V groups were exposed to the same perturbation a second time but under the other modality (auditory and visual, respectively). That is, participants who experienced the first perturbation under auditory (resp. visual) feedback received visual (resp. auditory) feedback during the relearning period. Participants of the V+A group received both modalities during the relearning period as well.

Surprisingly, in this second learning period, adaptation to the perturbation was observed under auditory feedback. Indeed, participants from the V→A group changed their reaching direction during the relearning period (change in angular error from start to end of relearning period: t_11_=3.4, p=0.006) even though they only received auditory feedback during that period. For this group, the difference between the learning under visual feedback and the relearning under auditory feedback did not reach significance (interaction between period of learning and time (early vs late): F_1,11_=4.62, p=0.05, 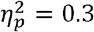). More importantly, this change in error was larger than the change observed during the first 100 trials of the learning period under auditory feedback in the A→V group (learning A→V group vs. relearning of the V→A group, interaction between group and time: F_1,21_=7.88, p=0.01, 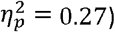).

The A→V group that received the auditory feedback during the learning period - and exhibited no learning - was well able to learn the visuomotor rotation under visual feedback (difference between initial and final errors in the relearning period: t_10_=4.2, p=0.002). This relearning was significantly larger than the change observed in the learning period of this group, under auditory feedback (interaction between period and time: F_1,10_=23.7, p=0.0006, 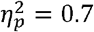). This suggests that this group is able to adapt to a visuomotor rotation under normal visual feedback condition but that the auditory feedback as provided here is insufficient to drive motor adaptation. Overall, we could not find any difference in the amount of relearning across the three groups (ANOVA on angular error, interaction between group and time: F_2,31_=0.35, p=0.7, 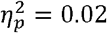).

### Relearning under auditory feedback is not due to an explicit strategy but to a memory of the learned directions

Two possible components of motor adaptation could account for the observed relearning under auditory feedback in the V→A group. Either participants adopted an explicit strategy and modified their reach angle consciously in the same direction as during the learning period, or they implicitly converged towards the learned reaching direction that was rewarded at the end of the learning period. To differentiate between these two hypotheses, we conducted an additional control experiment (Experiment 2) directly inspired by the experiment 3 of Huang et al (2011). In our experimental design, rotation angles between the learning (−25deg) and relearning periods (+25deg) are reversed. Therefore, the use of an explicit strategy would lead to an increase in error early during the relearning period. In contrast, the targets are arranged in such a way that the hand directions that counteract the visuomotor rotation are identical during the two adaptation periods. Therefore, the hand direction learned during the first adaptation period could drive the learning in the second adaptation period through reward-based learning even though the change in hand direction is opposite in these two periods.

**Figure 3.**
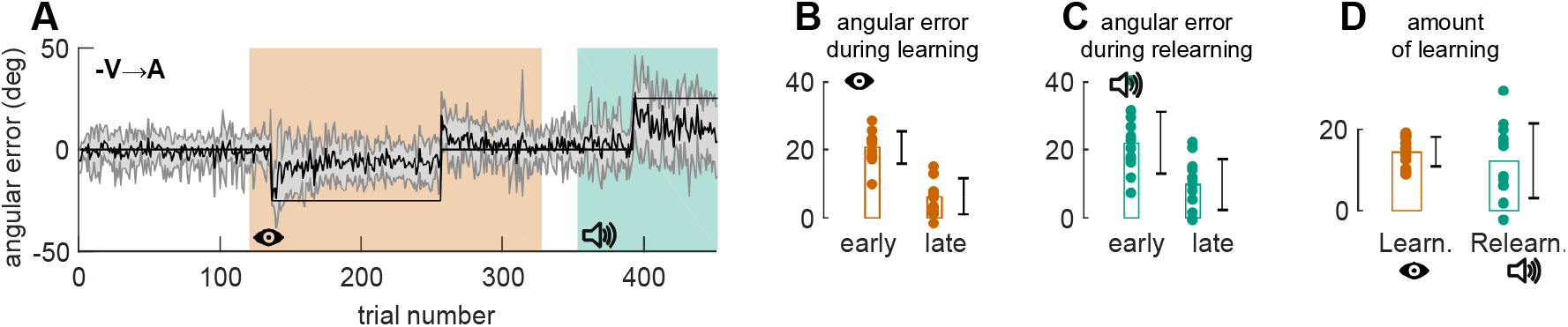
Evolution of the angular error over trials for Exp. 2. **A)** Evolution of the angular error over the course of the experiment. Data are averaged across participants (solid line) and grey shaded area correspond to the standard error of the mean. Thin black line corresponds to cursor rotation. Green background: auditory feedback. Red background: Visual feedback. **B)** Average angular error early and late during the learning period. Early = first cycle (three trials); Late=last six cycles (18 trials). **C)** Average angular error early and late during relearning. For **B, C and D**, colour code is similar to A. On each plot, individual values are represented by dots for each measure. Rectangles represent the mean. Black error bar represents mean ± SE.

Participants from this group (-V→A) adapted first to a −25deg visuomotor rotation under visual feedback (difference between early and late angular error, t_11_=13.74, p<10^−7^), which yielded a significant after-effect (mean ± SD: 12.2±6.2; t_11_=6.8, p<10^−4^). Most importantly, we observed a decrease in error during the relearning period (mean ± SD:12.3±9.1; t_11_=4.66, p=0.0007), consistent with the hypothesis that the reward-based memory of hand direction was able to guide the learning during the second adaptation period and that the explicit strategy was not responsible for the observed learning under auditory feedback during the second adaptation period.

The use of explicit strategy in the control experiment could be seen in an increase in the angular error during the first cycles of the relearning periods. However, such change in strategy was not apparent in our data as shown on Figure 4.

**Figure 4.**
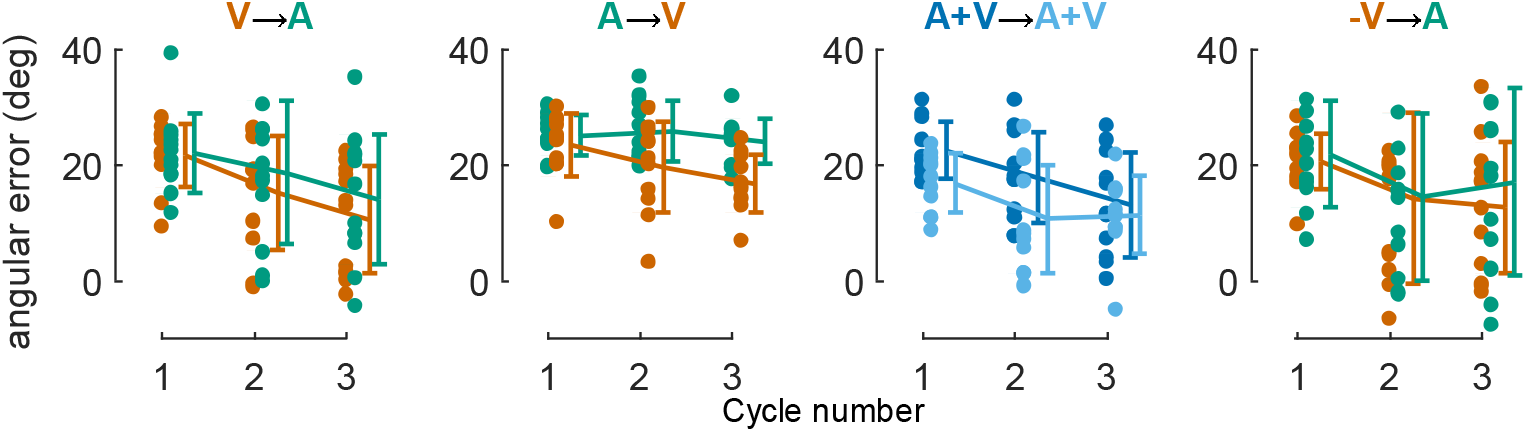
Evolution of the angular error during the first three cycles of the learning and relearning periods for the four experimental groups.

## Discussion

In the present study, we demonstrated that substituting visual error information with a auditory error information did not allow the participants to adapt to a 25deg visuomotor rotation. However, if adaptation to the perturbation by means of auditory error information was preceded by a first exposure to the perturbation combined with visual error information, then participants were able to adapt to the perturbation on the basis of auditory error information. Our control experiment suggests that adaptation under auditory feedback in the relearning period might be driven by a reward-based memory of the rewarded hand direction and not by an explicit strategy. Together, these results suggest that transfer of adaptation under different sensory feedback modalities is possible through reward-based motor memory that can act as an attractor for the reaching direction.

### Absence of learning on the basis of auditory error information on the first exposure

Humans are particularly skilled at detecting differences in pitch frequency. All our participants played at least one musical instrument. However, none was able to learn from auditory feedback when they received it during the first exposure. This finding is consistent with a previous study that showed that aftereffects were absent in auditory retention trials, whereas aftereffects remained high during visual retention trials (Kagerer and Contreras-Vidal, 2009). However, this also contrasts with previous studies (Oscari et al., 2012; Zanotto et al., 2013; Schmitz and Bock, 2014). These differences might be due to the experimental setup and to the way the feedback was provided as these are very different between our study and the previous ones. First, we provided endpoint auditory feedback while previous studies used online auditory feedback. It is known that online and endpoint feedback influence motor adaptation in different ways (Taylor et al., 2012). Second, we used the content of the auditory stimulus rather than its localization to provide feedback (Schmitz and Bock, 2014). This forces the participants to build an arbitrary mapping between the content of the auditory stimulus and the error they performed. This new relationship is very common when we cope with visuomotor rotations. For instance, moving a computer mouse on a horizontal table induces cursor displacements in the vertical plane (screen) with a gain usually different from 1. Third, there was a short delay between the end of the movement and the presentation of the auditory feedback, which might hamper the adaptation process (Brudner et al., 2016). This delay was the same in the VISUAL and AUDITORY modalities.

The absence of adaptation when auditory feedback alone was provided suggests that implicit adaptation of the internal model (Mazzoni and Krakauer, 2006; Morehead et al., 2017; Kim et al., 2018) might not take place under auditory feedback alone, as suggested by another study (Kagerer and Contreras-Vidal, 2009).

### Reward-based memory acts as an attractor

Interestingly, we found that, under auditory feedback, participants adapted to the perturbation during the second exposure but not during the first exposure. We believe that the learning on the basis of auditory feedback during the second exposure is driven by the spontaneous recovery of the motor memory (here a movement direction) elicited by the absence of reward (Pekny et al., 2011), which act as an attractor for the hand movement (Shmuelof et al., 2012). It is unsure whether the content of the auditory feedback played any role in the observed phenomenon. Indeed, given that the A→V group demonstrated that participants were unable to use the auditory feedback to adjust their hand movement to the visuomotor rotation, the observed effect might be driven by the removal of the reward at the end of each movement. In our experiment, successful movements were rewarded by points. Removing this rewarding signal yield to spontaneous recovery of the motor memory as observed in our study and in others (Pekny et al., 2011; Shmuelof et al., 2012). This is also consistent with the savings observed in experiment 3 of Huang et al. (Huang et al., 2011) and in Orban de Xivry et al. (Orban de Xivry and Lefèvre, 2015). Indeed, our control experiment that adopted a design similar to Huang et al. (Huang et al., 2011) shows that the sign of the error during the first exposure did not matter and that the learning during the second exposure was due to the recall of the reward-based memory, which acted as an attractor for the hand movement. The learning in the second exposure on the basis of the auditory feedback cannot be due to a memory of errors (Herzfeld et al., 2014) as the errors are radically different with the visual and auditory feedback. It is also unlikely that it is due to an explicit strategy as the explicit strategy would have driven the hand in the opposite direction in the control experiment (Morehead et al., 2015), which was not observed in the present study.

## Conclusion

In this study, we tested for the first time the independence of a reward-based motor memory of movement direction on the sensory modality during which it was created and found that reward-based memory can be formed and recalled independently of the sensory modality. This reward-based memory allowed the participants to learn a task that they could not learn otherwise independently of any memory of errors or explicit strategy. We believe that this study presents a new way of testing the presence and strength of a reward-based motor memory.

## Acknowledgments

This research was supported by the « Institut National de la Santé et de la Recherche Médicale » (INSERM), the « Conseil Général de Bourgogne » (France) and the « Fonds européen de développement régional » (FEDER).

